# DiNeR: a *Di*fferential Graphical Model for analysis of co-regulation *Ne*twork *R*ewiring

**DOI:** 10.1101/2020.05.29.124164

**Authors:** Jing Zhang, Jason Liu, Donghoon Lee, Shaoke Lou, Zhanlin Chen, Gamze Gürsoy, Mark Gerstein

## Abstract

**Background:** During transcription, numerous transcription factors (TFs) bind to targets in a highly coordinated manner to control the gene expression. Alterations in groups of TF-binding profiles (i.e. “co-binding changes”) can affect the co-regulating associations between TFs (i.e. “rewiring the co-regulator network”). This, in turn, can potentially drive downstream expression changes, phenotypic variation, and even disease. However, quantification of co-regulatory network rewiring has not been comprehensively studied.

**Methods:** To address this, we propose DiNeR, a computational method to directly construct a differential TF co-regulation network from paired disease-to-normal ChIP-seq data. Specifically, DiNeR uses a graphical model to capture the gained and lost edges in the co-regulation network. Then, it adopts a stability-based, sparsity-tuning criterion -- by sub-sampling the complete binding profiles to remove spurious edges -- to report only significant co-regulation alterations. Finally, DiNeR highlights hubs in the resultant differential network as key TFs associated with disease.

**Results:** We assembled genome-wide binding profiles of 104 TFs in the K562 and GM12878 cell lines, which loosely model the transition between normal and cancerous states in chronic myeloid leukemia (CML). In total, we identified 351 significantly altered TF co-regulation pairs. In particular, we found that the co-binding of the tumor suppressor BRCA1 and RNA polymerase II, a well-known transcriptional pair in healthy cells, was disrupted in tumors. Thus, DiNeR successfully extracted hub regulators and discovered well-known risk genes.

**Conclusions:** Our method DiNeR makes it possible to quantify changes in co-regulatory networks and identify alterations to TF co-binding patterns, highlighting key disease regulators. Our method DiNeR makes it possible to quantify changes in co-regulatory networks and identify alterations to TF co-binding patterns, highlighting key disease regulators.

## 1 Background

Thousands of transcription factors (TFs), their cofactors, and chromatin remodelers are employed in a highly coordinated manner to accurately initiate and control the transcriptional process on DNA sequences [1–4]. Precise temporal and spatial coordination among these factors is important for determining cell phenotype and maintaining biological function. Studies have reported that disruption of the co-regulation relationships of TFs can result in gene expression alterations, which can consequently introduce phenotypical variations and even lead to disease [5–9]. However, various computational methods have been proposed to infer dysregulations of an individual TF by exploring differential expressions of the TF itself and its gene targets [10], or investigating the direct TF-gene gain and loss events [11], ignoring the higher-order combinatory binding patterns among TFs. Therefore, large-scale mining of TF co-binding changes in disease and normal states could provide new insights into gene dysregulation and opportunities for targeted therapies.

In this study, we aimed to quantify such alterations to the TF co-regulatory relationships and prioritize regulators associated with pathogenesis. This is a challenging task for many reasons. For instance, many of the ~1,400 known human TFs have highly dynamic binding profiles depending on the cell state and conditions [12]. Therefore, it is essential to carefully curate a dataset of appropriate cell types to precisely capture disease-specific TF co-regulatory disruptions. Furthermore, joint analysis of binding profiles of numerous factors would be beneficial by maximizing our knowledge of TF co-binding events. Researchers have proposed various models to systematically impute all tissue-specific TF regulomes using features such as DNase accessibility and sequence context, but the accuracy of these methods across TFs is still unknown [13, 14].

In light of this, we propose a multi-step computational framework that we call <u>*Di*</u>fferential graphical model of <u>*Ne*</u>twork <u>*R*</u>ewiring (DiNeR) to infer TF co-binding alterations and pinpoint disease-causing TFs. First, we modeled the cooperative regulation patterns among TFs using a co-regulation network, where the nodes represent TFs and the weighted edges measure the genome-wide co-occurrence between pairwise TFs, all of which are derived from chromatin immunoprecipitation followed by sequencing (ChIP-seq) data. This is distinct from traditional regulatory networks, where edges usually imply the relationship between a TF and the physical interaction it shares with an enhancer or promoter region in order to initiate the transcription process of its target genes. Second, we directly measured the gain or loss of TF co-regulatory events across the genome during the transition from one cellular state to another using a differential graphical model. Intuitively, a higher weight in our differential network indicates a larger alteration in the co-regulatory pattern between two states. As a third step, we adopted the graphical LASSO model and used a stability-based method for penalizing parameter selection to control for the sparsity of the differential network with the goal of removing spurious edges [15–18]. Lastly, we prioritized the TFs according to their degree in the differential network based on the assumption that disease-driving TFs can demonstrate massive binding profile changes and consequently disrupt their cooperative regulation with many other regulators.

To test the effectiveness of our DiNeR framework, we applied this model to the ENCODE Tier 1 cell lines, K562 and GM12878, roughly representing paired disease-to-normal cells for chronic myelogenous leukemia (CML). We used our DiNeR framework to directly estimate the changes in the TF combinatory regulation relationships between disease and normal states. As a result, we identified several TFs that exhibit a high level of disruption to their co-binding partners. Among this list were several well-known risk factors for leukemia, such as BRCA1, RAD51, BMI1, and H3k27me3 [19–22], demonstrating the effectiveness of our method.

## 2 Results

### 2.1 Collecting appropriate large-scale ChIP-seq data to investigate transcriptional regulation

TF binding sites are highly tissue specific and usually change dramatically across cell types. Therefore, it is important to select appropriately matched and representative cell types to investigate a specific disease. Here, we used K562 and GM12878 to represent a rough disease-to-normal pair in order to investigate the transcriptional regulation dynamics during oncogenesis for CML (see details in section 5.1). Specifically, we extracted 94 common TFs among these two cell lines (Fig. 1A, Table S1). In order to investigate alterations in the joint activity between TFs and specific chromatin marks in disease, we also extracted nine histone modification marks and chromatin accessibility data sets from these cell lines. Among these TFs, 31 showed significant expression changes between disease and normal states (Fig. 1C).

**Figure 1.**
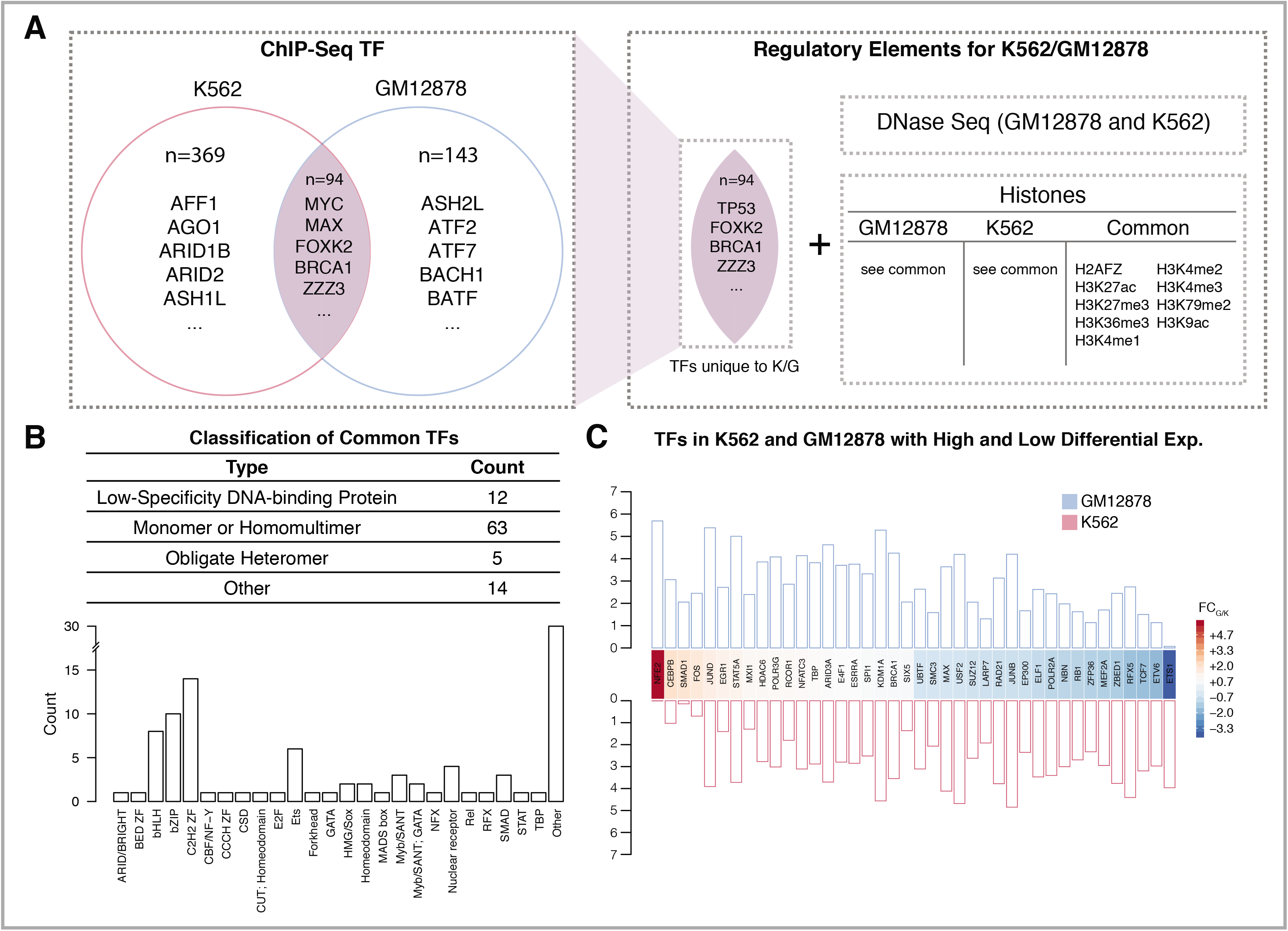
ChIP-seq data collection and pre-processing. A) TFs shared by both the K562 (red) and GM12878 (blue) cell lines from ENCODE, in addition to DNase sequencing and histone modification data, are pooled together to form the set of regulatory elements used in the analysis. B) Classification of the common TFs between K562 and GM12878. Specific DBD family classifications are also given. C) The heatmap shows the top and bottom 20 TFs with high and low log fold change of expression between GM12878 and K562. The barplot shows the log of the FPKM expression values for each of these TFs.

### 2.2 Building a genome-wide TF co-regulatory network

Aberrant transcriptional regulation is associated with various diseases [5–9]. Network analysis has been proven to be a powerful tool for identifying and prioritizing genes or regulators associated with pathogenesis. For example, scientists have constructed TF regulatory networks in order to mimic the physical binding of TFs to either enhancer or promoter regions during initiation of the transcriptional process of its target gene (Fig. 2A, B) [11]. Edges in this type of regulatory network only focus on the local interaction between the TF and target gene pair, and do not consider the effect of the rest of the genome. Another approach is to use gene coexpression networks, where a shared edge in the network represents consistent expression patterns between a pair of genes across many samples, which was usually inferred from RNA sequencing or microarray data [23, 24]. However, this output only reflects co-expression patterns, rather than regulatory relationships.

**Figure 2.**
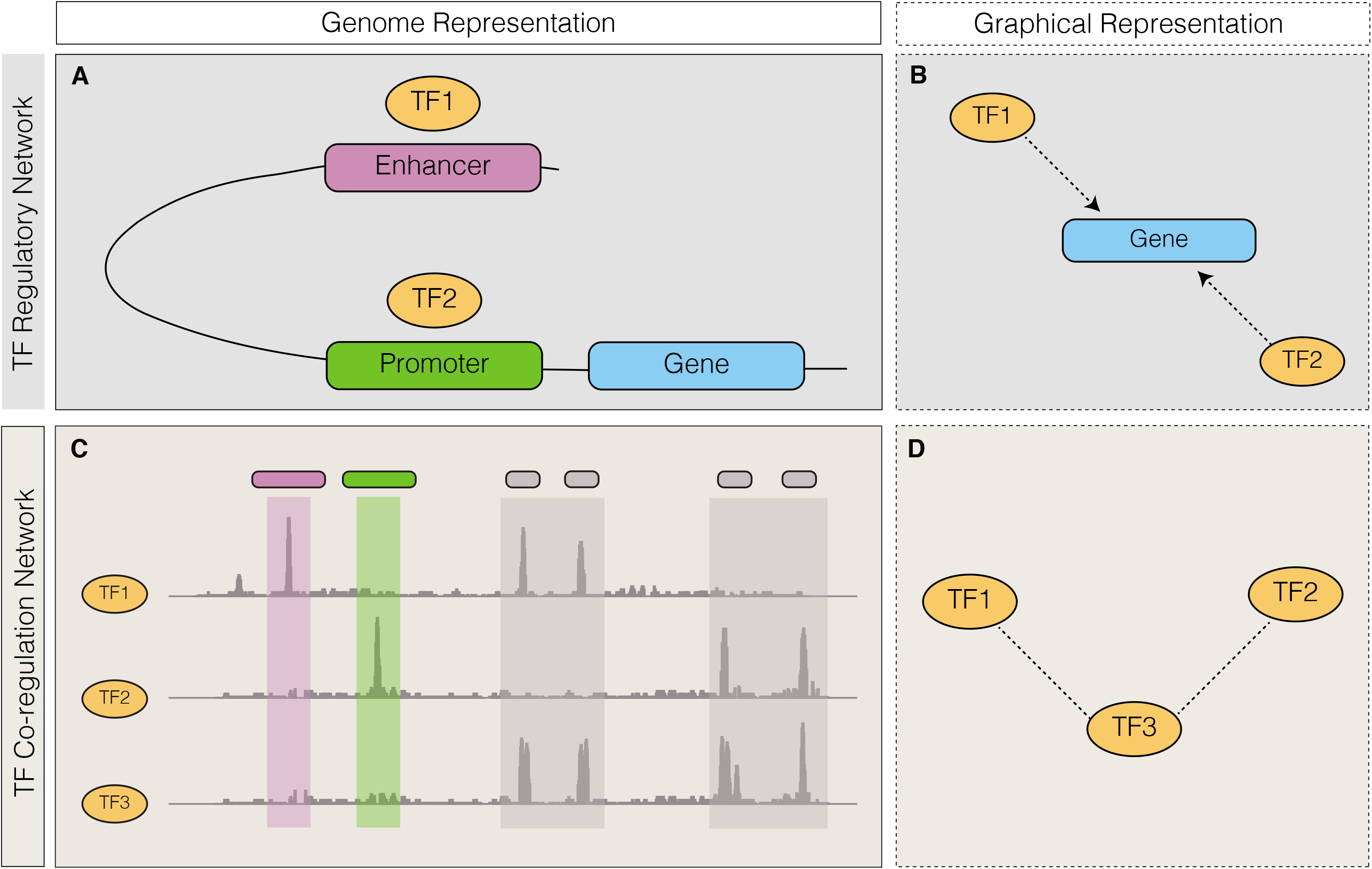
A schematic of two types of networks are given, in both genomic and graphical form. A) A regulatory network is given in genomic form where TF1 and TF2 both regulate the downstream gene, binding to an interacting enhancer and promoter, respectively. B) A directed graph is shown where TF1 and TF2 both regulate the gene. C) A genomic representation of a co-regulatory network is shown, based on the signal tracks of three TFs. D) A graphical form of the co-regulatory network is shown, where TF3 co-binds with TF1 and TF2 separately.

Here, we propose a TF co-regulatory network based on large-scale ChIP-seq data to model a related but different aspect of transcriptional regulation – the cooperative behavior among TFs. Specifically, in our network, nodes represent TFs and the weighted edge between them measures the level of non-random co-binding activity across the genome. It is important to note that this network is distinct from a traditional TF-TF network, which focuses on the mechanism of how one TF gene is regulated by another TF protein. In particular, this type of TF-TF regulatory network describes the binding event of one TF at the non-coding element (e.g., enhancer or promoter) of another TF, but does not indicate whether these two TFs work in a coordinated way to regulate other downstream genes.

We used the Gaussian graphical model (GGM) to construct this TF co-regulatory network [25]. The schematic in Fig. 2C illustrates one example. Two pairs TF_1_ and TF_3_, TF_2_ and TF_3_ co-bind over many places in the genome, possibly resulting from interacting domains that are necessary to initiate transcription in many genes. Then we draw two edges in our co-regulation network between TF_1_ and TF_3_, TF_2_ and TF_3_ (Fig. 2D).

### 2.3 Using differential networks to measure TF co-regulation alterations between states

After building the co-regulatory networks under one condition, we aimed to provide a quantitative measure of co-regulation alterations between two conditions (e.g., disease and normal states). Intuitively, our goal is to try and infer a differential co-regulatory network. In this differential network, while nodes still represent TFs, each edge now describes the level of change that each TF pair experiences during disease progression (Fig. 3). Therefore, we extended the GGM for one condition to a differential graphical model for two conditions by estimating the differences between two precision matrices (details in section 5.2.2).

**Figure 3.**
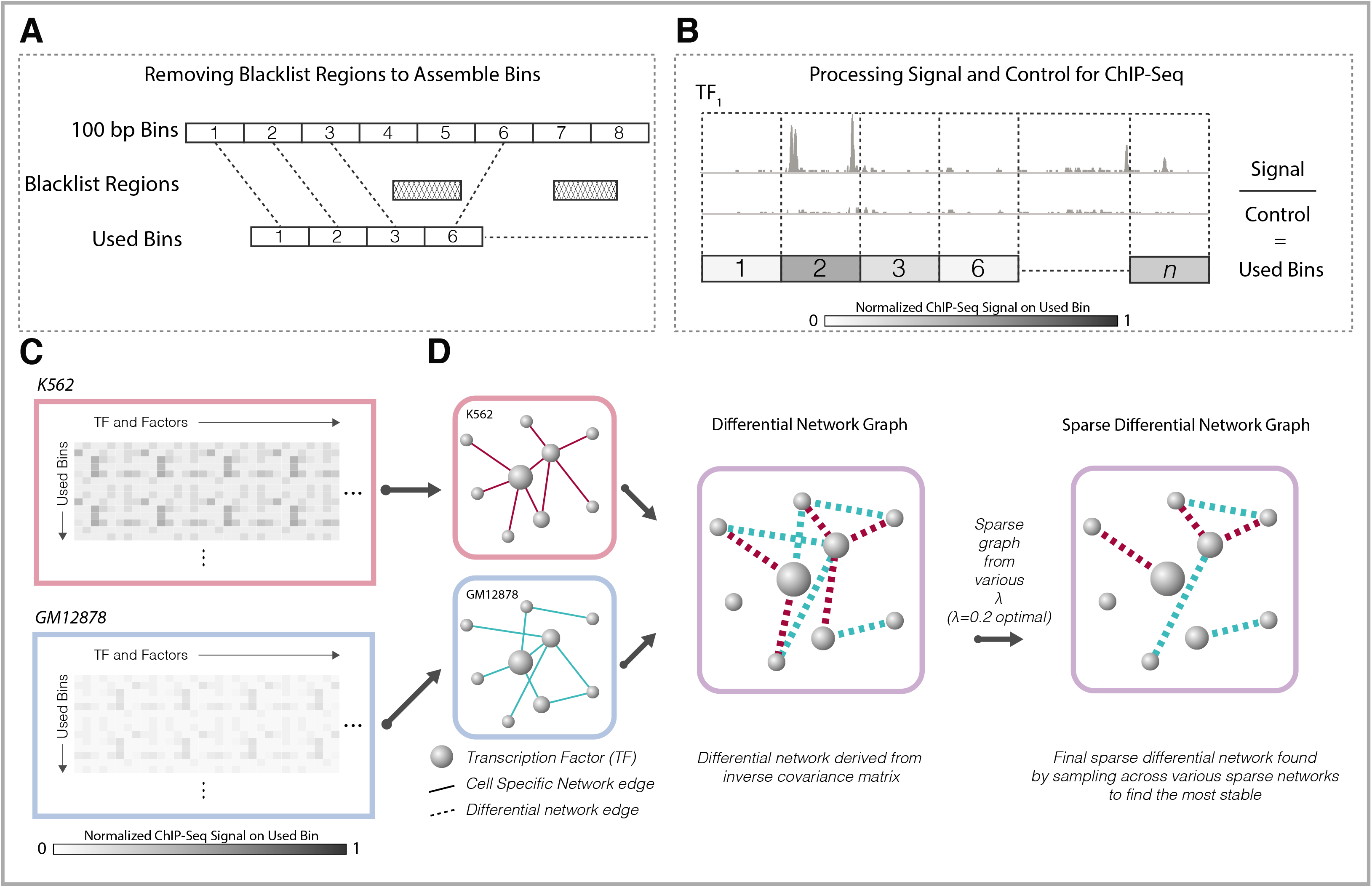
Workflow of DiNeR method to create sparse differential regulatory network. A) Bins used for the analysis are created by taking 100bp bins on the genome and removing any blacklist regions to form the used bins; B) For each TF and factor, the ChIP-Seq signal fold change is averaged over the used bins; C) For disease and control cell types K562 and GM12878, respectively, a fold change signal matrix is created with rows or bins and columns of TFs and factors; 4) By using difference of the precision matrix, a differential network is created, which is then made sparse through sampling-based penalization.

During the estimation process of these two precision matrices, small non-zero values are introduced and may lead to many spurious edges, resulting in a dense differential network with many false positives. Therefore, we introduced a regulation parameter, *λ*, to penalize the retained edge count in order to remove potential spurious edges. As shown in Fig. 4, higher *λ* values indicate a larger penalty in the network edge, resulting in a sparser estimation of the differential network.

**Figure 4.**
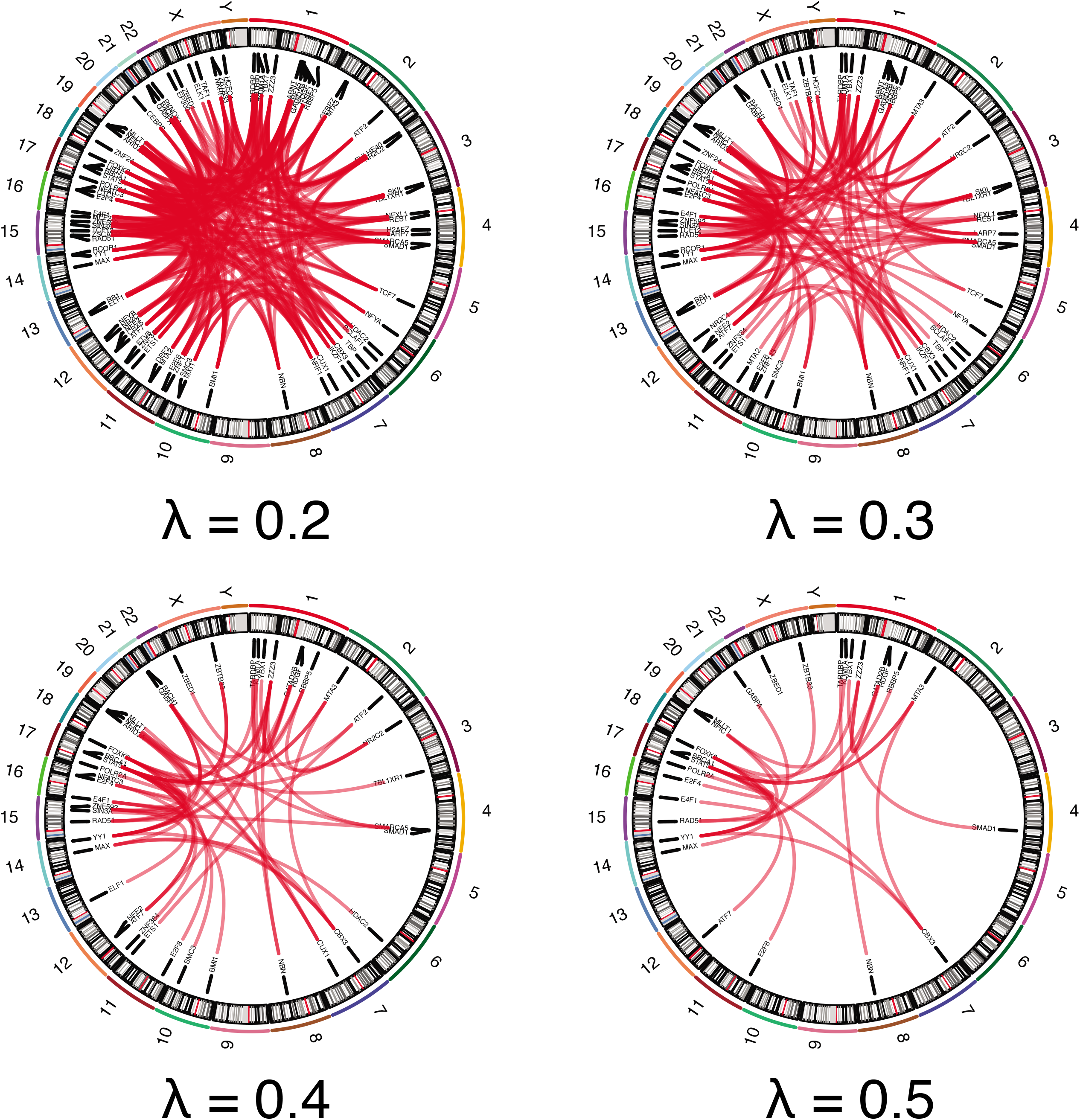
Circos plots are used to show the sparsity of the network created using various lambda values. λ values of 0.2, 0.3, 0.4, and 0.5 are used and their networks are shown.

Selecting an appropriate *λ* parameter is key to reliably inferring the TF co-regulation gain and loss events while simultaneously removing false positives. We further used a stability-based model selection method to choose the optimal *λ* based on sub-sampling of the genome [15]. The intuition of our model is that we encourage the network to be more inclusive of edges for the benefit of allowing us to scrutinize many possible changes associated with disease, while simultaneously ensuring that the results are highly repeatable across many regions in the genome (Fig. 5, details in section 5.3).

**Figure 5.**
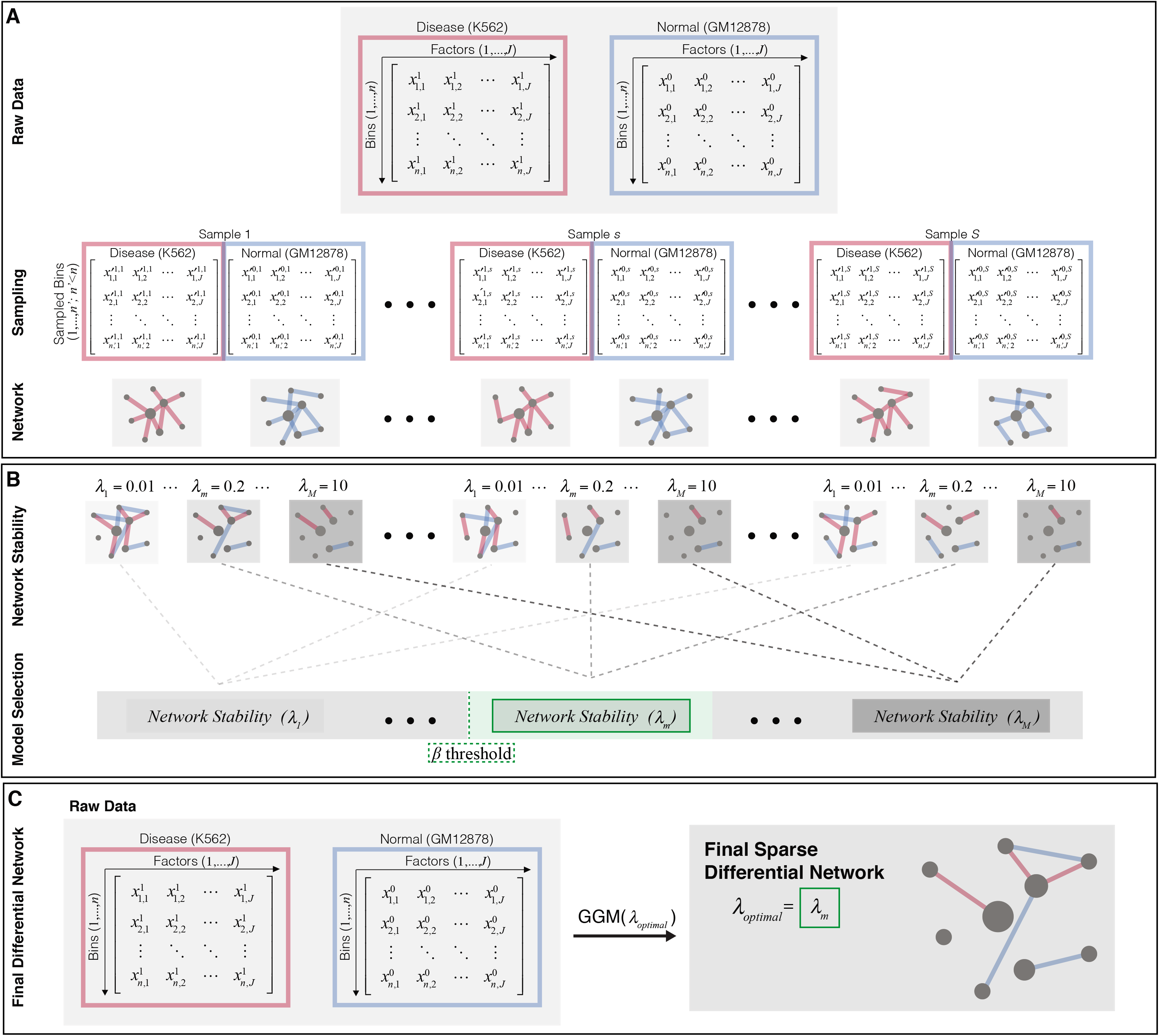
A flowchart of the model selection process to create a sparse differential network is shown. A) Raw data is sampled and graphical representations of the network are given for disease and control. The samples are different and therefore have varied network representations. B) Differential networks are generated using GGM using various *λ* parameters, *λ*:{1,…, *λ_M_*}, and a model selection is done through stability analysis to see which *λ* gives the most stable network variance. A threshold for stability, *β*, allows us to choose the optimal *λ*, shown in green. C) The optimal *λ*, in this case *λ_m_* is used on the original raw data, to generate the final sparse differential network.

### 2.4 Applying DiNeR to prioritize key TFs associated with CML

We applied our DiNeR framework to 104 paired factors in K562 and GM12878 cell lines. After model selection, we set *λ_opt_* − 0.2 (details in section 5.3). In total, we included 6.49% of all possible edges in the final network (351 out of 5,408). We identified eight out of the 104 factors as consistent network hubs (Table 1), and reliably captured all of them by sub-sampling half of the genome. These eight factors include many well-known genes that have been previously associated with leukemia (Table 1), indicating that our method can reliably detect key regulators of disease.

**Table 1:**
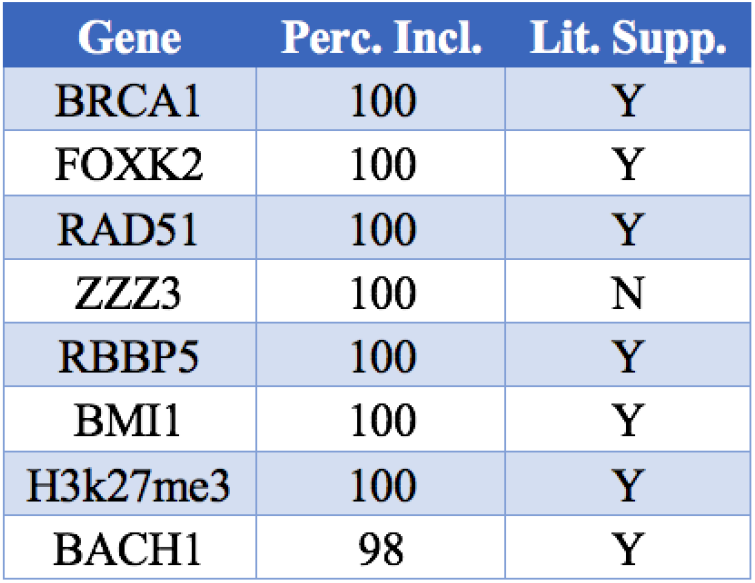
A list of DiNeR prioritized TFs using the network hubs of the K562 vs. GM12878 differential co-regulation networks upon different subsampling of the entire genome. Perc. Incl.: how many rounds of simulations out of 100 this TF has been claimed as a network hub. Lit. Supp: whether there is literature support to link this TF with cancer.

| Gene | Perc. Incl. | Lit. Supp. |
| --- | --- | --- |
| BRCA1 | 100 | Y |
| FOXK2 | 100 | Y |
| RAD51 | 100 | Y |
| ZZZ3 | 100 | N |
| RBBP5 | 100 | Y |
| BMI1 | 100 | Y |
| H3k27me3 | 100 | Y |
| BACH1 | 98 | Y |

One of the identified factors was the proto-oncogene BMI1, which is a major component of polycomb group complex 1 and plays a central role in DNA damage repair. Although BMI1 showed only moderate expression changes in tumor as compared to normal cells (approximately 20% higher; from 42.90 to 51.19), its co-binding relationship with other factors changed dramatically. BMI1 was a significantly rewired TF and maintained edges with 14 other factors, including a well-known cancer-related histone modification mark H3K27me3. Interestingly, BMI1 is a known biomarker of hematologic malignancies and has been shown to be essential for faithful reprogramming of myeloid progenitors [26, 27]. Studies have reported that levels of BMI1 correlate with prognosis of patients with myelodysplastic syndrome and chronic and acute myelogenous leukemia [21]. Our analysis provides another possible explanation to this phenomenon, suggesting that in addition to aberrant expression patterns, the disruption of the coordination between key regulators during transcription can also contribute to disease progression.

Another interesting factor identified by our method was the tumor suppressor gene BRCA1 (Fig. 6A). BRCA1 has been shown to play a central role in maintaining genomic stability. Moreover, germline defects in this gene have been associated with multiple cancer types, including breast and ovarian cancers [28]. In normal cells, BRCA1 is typically associated with RNA polymerase II across multiple species to faithfully activate transcription of key genes [29]. However, we found that in K562 cells, the key interaction between BRCA1 and RNA polymerase II was significantly disrupted. Specifically, the Jaccard distance between the BRCA1 and POLR2A ChIP-seq peaks decreased by 98% in K562 as compared to GM12878 cells, indicating severe colocalization disruption. Consistent with this finding, we observed a remarkable proximal-to-distal shift in the BRCA1 binding locations in K562 cells (Fig. 5AB). In particular, approximately 93.6% of the BRCA1 ChIP-seq peaks in GM12878 were located within a 5 kb region of annotated transcription start sites (proximal), and this number was drastically reduced to 16.4% for K562 peaks. Such a remarkable shift indicates that in cancer cells, BRCA1 fails to cooperate with other conserved collaborators to control the transcriptional process. BRCA1 has also been widely reported to affect chromatin structure and introduce chromatin remodeling. Interestingly, we also found that H3K4me1 and open chromatin regions were among the rewired partners of BRCA1. The overlapping peaks between BRCA1 and these factors were significantly reduced during the normal-to-tumor transition (p value < 2 × 10^−16^ for binomial tests in both cases). Hence, we hypothesize that BRCA1 severely alters not only its transcriptional regulation through promoter region interactions, but also its chromatin remodeling activity.

**Figure 6.**
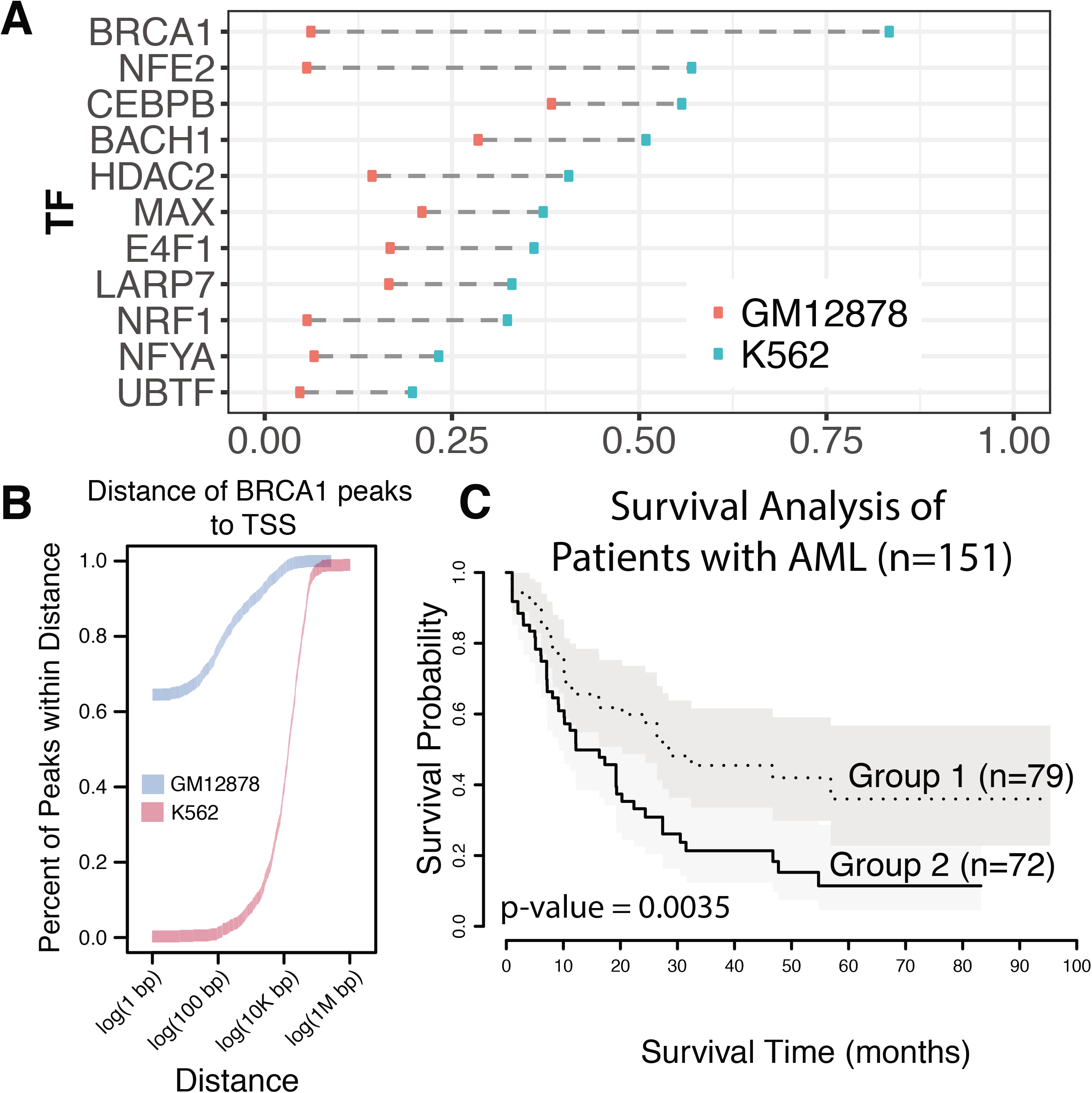
A) Summary of the proximal to distal change of binding peaks from GM12878 to K562. B) Cumulative density of distance of BRCA1 peaks to the nearest TSS in both GM12878 and K562 showcases a dramatic shift in binding profiles between disease and control. C) Survival analysis in AML using BRCA1 regulatory activity as a prognosis marker, showcasing disrupted BRCA1 regulatory activity results in a worse prognosis.

### 2.5 Further investigating TFs prioritized by DiNeR

To further explore the prognostic value of BRCA1 in leukemia and validate our model, we downloaded 151 patient expression and clinical profiles from The Cancer Genome Atlas (TCGA) for acute myeloid leukemia [30]. If we only consider the expression levels of BRCA1 in K562 and GM12878, we are unable to fully detect the difference in behavior of this gene between the two cell types. However, when considering the regulatory activity, we found that BRCA1 was upregulated by 50% in K562 as compared to GM12878 cells. In line with large-scale network rewiring events, we combined the BRCA1 regulatory network with patient tumor-to-normal differential expression profiles to measure the regulatory potential of BRCA1 in each patient. In contrast to the survival analysis using expression profiles alone, we found that a severe disruption of BRCA1 regulatory activities usually indicated worse patient survival rates (P value = 0.0035) (Fig. 6C). This analysis demonstrates that the differential graphical model can effectively identify key factors affecting patient survival beyond gene expression.

## 3 Discussion

Simultaneous binding of many TFs to proximal and distal regulatory regions of the genome is imperative for precise control of spatiotemporal gene expression patterns. Therefore, it is essential to investigate transcriptional regulation alterations in order to prioritize risk factors associated with disease. Many methods have been developed to address this goal. For instance, researchers investigated TF target gene gain and loss events and found distinct patterns between oncogenes and tumor suppressors [11]. Others combined expression changes with TF regulatory networks and identified TFs responsible for aberrant gene expression patterns, and then validated their discoveries through patient survival analysis [10]. Here, we focused on a unique aspect of transcriptional regulation – coordination among TFs, which goes beyond the binding or expression changes of TFs. For instance, genome-wide chromatin remodeling and histone modifications changes may introduce dramatic binding profile alterations for different TF, resulting in alterations of the combinatory coordination among TFs. For instance, some TFs, such as BRCA1 in our analysis, demonstrate massive binding profile changes. As a result, these TFs disrupt the co-binding relationship with many other TFs and chromatin features and are prioritized with the highest importance by our DiNeR framework. We also found that the target genes of TFs showed distinct expression patterns in tumors compared to normal cell lines, most likely due to extensive binding profiles changes. As a result, our analyses offer related but complementary views of transcriptional dynamics compared to previous methods and provide new insights into disease-related transcriptional regulation.

We also emphasize that model selection is a key step of our DiNeR framework. A larger penalty to the edge numbers captures the more confident co-regulatory network alterations at the cost of excluding weaker, but not necessarily insignificant, changes. Researchers have proposed various model selection methods, such as Akaike and Bayesian information criteria, to solve this problem [25]. In our framework, we used the stability-based model selection method, which provides a more interpretable explanation [15]. Specifically, we include an edge in the network as long as we can reliably discover it from numerous random subsamples of different genomic regions.

In addition, we ranked TFs according to the number of gained and lost partner TFs in the differential network. The intuition behind this scheme is that TFs that show a larger degree of genome-wide binding profile changes represent network hubs and are more likely to have disrupted coordination with many other partner TFs, resulting in a higher impact on transcriptional regulation. This is a reasonable assumption without any prior information. However, it is possible that even the disruption of one canonical TF pair may change the expression of key genes, such as in cases of oncogenes and tumor suppressor genes. Hence, it is also valuable to scrutinize the non-hubs of the network provided by DiNeR.

Finally, we showed that using matched ChIP-seq data from disease and normal cells is beneficial to directly capturing the co-localization events of TFs, as compared to other profiles such as expression. However, this approach requires the availability of hundreds of functional characterization datasets. We believe that as high-throughput sequencing technologies continue to develop, especially single-cell sequencing methods, our proposed differential graphical models could be applied to new opportunities to highlight regulators in disease.

## 4 Conclusion

We developed a TF-TF network rewiring and regulator prioritization method by applying nonparametric graphical models on large-scale functional genomics data. This approach allows us to identify DNA binding factors demonstrating large co-localization disruptions. Given the number of genome-wide binding profiles from ChIP-seq data in matched disease and control cells, our DiNeR method can reliably and efficiently highlight the significant changes of pairwise coordinated regulations between different factors. We applied our model to 104 common TF, histone modification, and chromatin accessibly data from a loosely paired tumor and normal cell line in CML. We discovered disruptions between well-known partners of transcriptional regulation, such as BRCA1 and RNA polymerase II, signifying the effectiveness of our method.

## 5 Methods

Here, we adopted a differential graphical model to investigate the differences in co-binding patterns of TFs between normal and disease conditions. We hypothesized that factors that dramatically change their partners during the transcription process would be altered to a larger degree in disease samples, and hence would have larger effects in driving disease progression.

### 5.1 ChIP-seq data collection and pre-processing

We aim to infer the differential TF co-regulation alterations among normal and disease conditions. Toward this goal, we searched for disease-to-normal TF pairs with at least 50 common TFs and found that only the K562-GM12878 pair satisfied this requirement. Therefore, we decided to use the 369 and 143 ChIP-seq experiments from K562 and GM12878 cell lines, respectively. After de-duplicating and extracting common ChIP-Seq targets, we identified 94 common TFs among these two cell lines (Table S1). In order to investigate alterations in the joint activity between TFs and specific chromatin marks in disease, we also extracted nine histone modifications and chromatin accessibility datasets from these cell lines (Fig. 1A,B). The majority of the 94 common TFs were sequence-specific binding factors (TFSS in Fig. 1C), 31 of which showed significant expression changes between disease and control.

To uniformly process the data, we first divided the autosomal chromosome (hg38 version) into 100 base pair (bp) bins and removed bins that overlapped with genomic regions that have gaps or low mappability using BEDTools (version 2.27.1-foss-2018b)[31]. To remove any artifacts from non-peak regions, we further removed all 100 bp bins that did not overlap with any peaks from these 104 factors (Fig. 3A). In total, we kept 1,351,140 bins in our analysis.

For each factor, we calculated the normalized read count for the ChIP-seq experiment versus its matched control experiment for all bins (Fig. 3B). We then calculated the average signal from each of the replicates or experiments from different labs when multiple datasets were present for a single factor. Details of the ChIP-seq signal files used has been listed in the supplementary data. We organized the resultant signal data into a matrix for K562 and GM12878 separately, with columns indicating factors and rows indicating bins in the genome (Fig. 3C). We used these two matrices as the inputs for the following differential graphical model (Fig. 3D).

### 5.2 Infer differential graphical model to TF-TF network rewiring

#### 5.2.1 Gaussian graphical model for co-regulation network in one state

In this section, we describe the details of the differential graphical model. Let *G*^(*D*)^ = (*V*^(*D*)^, *E*^(*D*)^) represent the network with nodes and edges *E*^(*D*)^ for disease status *D*. Let 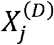 denote the vector of the average normalized ChIP-seq signal of factor *j* in status *D*, where *j* = 1… *J. D* = 1 indicates disease and 0 indicates normal. Next, for each *D*, we calculated 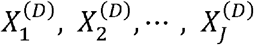 over 100 bp bins for all the 104 factors. In our analysis, we included 94 TFs, nine histone modification marks, and one chromatin accessibility (*J* = 104).

Under one condition *D*, we assume that 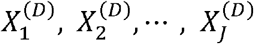 follows a multivariate Gaussian distribution such that 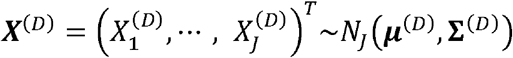. We can construct the TF co-regulatory network using a traditional GGM. Here, we are aiming to identify TF true physical interactions by highlighting conditional dependent binding profiles among TFs. Therefore, we used the precision matrix represented as **Θ**^(*D*)^:= (**∑**^(*D*)^)^−1^ to infer whether any TF pair has a non-random co-binding interaction. In other words, If 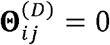, then 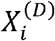 and 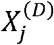 are independent of each other, conditioned on all the other TFs. As a result, 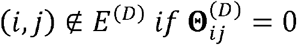.

#### 5.2.2 Differential graphical model for co-regulation alteration in two states

Next, we used the difference between two networks *G*^(1)^ and *G*^(0)^ called the differential network, to represent the degree of TF co-regulation alteration under two conditions (*D* = 1 and *D* = 0). Given the observed data 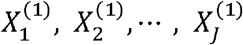 in the disease cell and 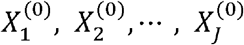 for the normal cell, edges in the differential co-regulatory network can be inferred from the difference between the two precision matrices **Δ** = **Θ**^(1)^ − **Θ**^(0)^ = (**∑**^(1)^)^−1^ − (**∑**^(0)^), where the co-regulation relationship between *TF_i_* and *TF_j_* changes if |Δ_*i,j*_| ≠ 0. Note that **∑**^(1)^**Δ∑**^(0)^ − (**∑**^(1)^ − **∑**^(0)^) = 0. Hence, we can solve the following equation to estimate a reasonable **Δ**.

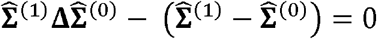

Here 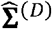 is the sample covariance matrix. Specifically, we used the penalized D-Trace loss model estimate **Δ** [16–18].

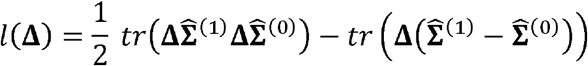

*tr* represents the trace of a matrix. To remove spurious differential edges, we introduced a nonnegative regularization parameter *λ* to penalize the number of edges in the network.

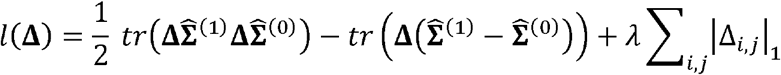

Here, *λ* controls the sparsity of the rewired network. For example, *λ* − 0 indicates no penalty and usually will result in a very dense network. In contrast, a large *λ* value will result in a sparse network.

#### 5.2.3 Co-variance matrix inference

Under the Gaussian assumption, 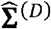 can be directly obtained from the sample covariance matrix. However, One statistical concern of using a differential graphical model is the distribution of **X**^(*D*)^. Here, we found that even after log transformation, almost all TFs severely contradicted the Gaussian assumption (for details see suppl. sect. S2). Therefore, going forward we used a nonparametric model instead of a GGM. Our assumption is that a set of monotonically increasing functions 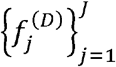 exists such that, after transformation, 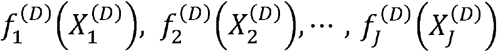 follow a multivariate normal distribution *N_J_*(**O,∑**^(*D*)^). Similar to the GGM, we can use the precision matrix *Θ*^(*D*)^: = (**∑**^(*D*)^)^−1^ to infer the conditional dependence between any pair of factors in the network. As described in [17], we adopted the rank-based scheme to estimate the sample covariance matrix without directly estimating 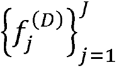. Specifically, let 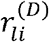 represent the rank of bin *I* for TF *i* in status *D* among all the bins, and *n* is the total number of bins in the genome. The Spearman correlation of TFs *i* and *j* are represented as below.

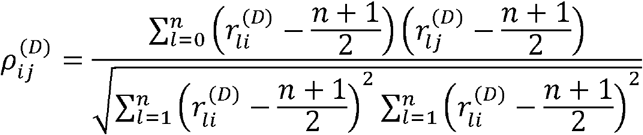

Then we replace the sample covariance matrix 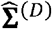 by, 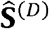 with elements

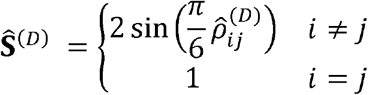

In cases where 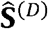 was not positively semi-definite, we used a projection method as described in [17, 32].

### 5.3 Model selection

It is critical to select an appropriate *λ* to reliably infer network changes in disease samples because different *λ* values can lead to different conclusions in downstream analyses. Researchers have proposed many methods, cross validation, Akaike information criterion (AIC), and Bayesian information criterion (BIC), to automatically select *λ* [33–35]. We chose a more interpretable approach in our ChIP-seq-based co-regulation network analysis than AIC and BIC, by using the Stability Approach to Regularization Selection (StARS) approach (Fig. 5) [15]. The key characteristic of this method is that it encourages the differential network to be inclusive to account for the true dynamics between disease and control networks, while guaranteeing an acceptable stability in the resultant differential network.

Specifically, we defined 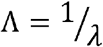 as an alternative parameter to control network density so that a larger Λ indicates a denser network. During the model section process, we start from subsampling part of the genome for S’ times. Specifically, during the *s^th^* sampling, ***X***^(1),*s*^ and ***X***^(0),*s*^ represent the binding profile matrices. 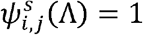 if there is an edge between TF *i* and TF *j* in the rewired network under Λ, otherwise 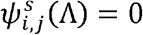. In the *s* = 1,2,…,*S* randomly sampled datasets, we defined 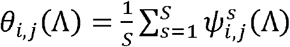, and *ξ_i,j_*(Λ) = 2*θ_i,j_*(Λ){1 − *θ_i,j_*(Λ)} to be the fraction of times the networks disagree with the existence of the edge (*i,j*). Then, the overall instability of the networks over the sampling sets is

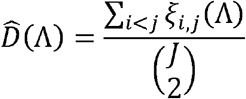

It is clear that 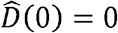 in an empty network because there is no instability when there are no edges. In general, the network becomes denser and more instable as Λ goes larger. However, when the network becomes very dense and even fully connected, 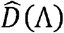 goes smaller again and eventually reduces to zero. As suggested in [15], we used the monotone function 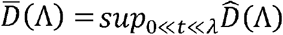 to remove such artifact effect in an extremely dense network. As a result, the optimal Λ should be 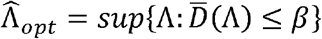 for a predefine network instability measure *β*.

In our analysis, we started from a broad spectrum of parameter values from *λ*_1_ = 0.01, *λ*_2_ = 0.05,…*λ_m_*,…,*λ_m_* = 10, representing a wide range of sparse networks, from almost fully connected to empty (Fig. 4). For each *λ_m_*, we randomly selected half of the bins we used in section 5.2 to run the LASSO penalized D-Trace loss model. We repeated this process *S* = 100 times for each *λ_m_* and calculated the average network variance. We used *β* = 0.5% as our stability threshold and selected the optional *λ* is 0.2.

## 6 List of Abbreviations

DiNeR: <u>*Di*</u>fferential Graphical Model of co-regulation <u>*Ne*</u>twork <u>*R*</u>ewiring to Infer Transcription Factor Co-binding Alterations
TF: Transcription factor
CML: chronic myeloid leukemia
GGM: Gaussian graphical model
ChIP-seq: Chromatin immunoprecipitation followed by sequencing
StARS: Stability Approach to Regularization Selection
TCGA: The Cancer Genome Atlas

## 7 Declarations

### Ethics approval and consent to participate

Not applicable

### Consent to publish

Not applicable

### Availability of data and materials

The datasets generated and/or analyzed during the current study are available in the ENCODE portal, https://www.encodeproject.org/ [21]. Detailed ChIP-seq data (with accession code) used in this work has been listed in detail in the supplementary data. Users can directly download the raw data used in this paper from the ENCODE website with the accession code.

### Authors’ contributions

JZ designed the methods; JZ, JL, DL, and ZC performed all the analyses; SL and GG helped with expression data processing; all authors were involved in writing the paper.

## Acknowledgements

We thank the Yale Center for Research Computing for guidance and use of the research computing infrastructure at Yale university.

