## Supplement for "DiNeR: a *Di*fferential Graphical Model for analysis of co-regulation *Ne*twork *R*ewiring"

Supplementary file to

"DiNeR: a nonparametric ***Di***fferential Graphical Model on ***Ne***twork ***R***ewiring to Infer Transcription Factor Co-binding Alterations in Disease"

#### S1. Rationale to match K562 to GM12878

Exact matching was not possible with K562: this cancer cell-line was derived from a myeloid lineage, but there is no data-rich, non-cancerous myeloid cell assayed in ENCODE. GM12878 is a data-rich ENCODE cell-line derived from the closely related lymphoid lineage. Supporting this choice, we determined that among all non-cancerous cell-lines provided by Roadmap Epigenome and GTEx, GM12878 has the highest Spearman correlation with K562 in gene expression. Hence, we used GM12878 as the most appropriate normal pair for K562.

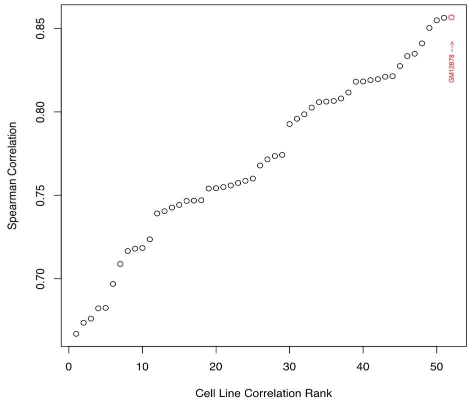


*Figure S 1. Expression matching between cell types*

#### S2. S3. Violation of normal assumption

We calculated the signal of each bin for all uniformly processed ChIP-seq data from ENCODE using bigWigAverageOverBed. Then, we took the log of the signal over the genome and plotted the density Figure S 2. We also plotted the QQ-plot vs. theoretical normal distribution, as shown in Figure S 3. The P-value of the Kolmogorov-Smirnov test for normality is less than $2.2e-16$, indicating a strong violation of the normal assumption.

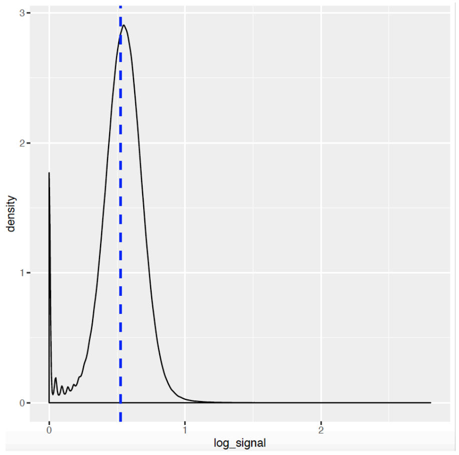

*Figure S 2. density plot of the ChIP-seq signals*

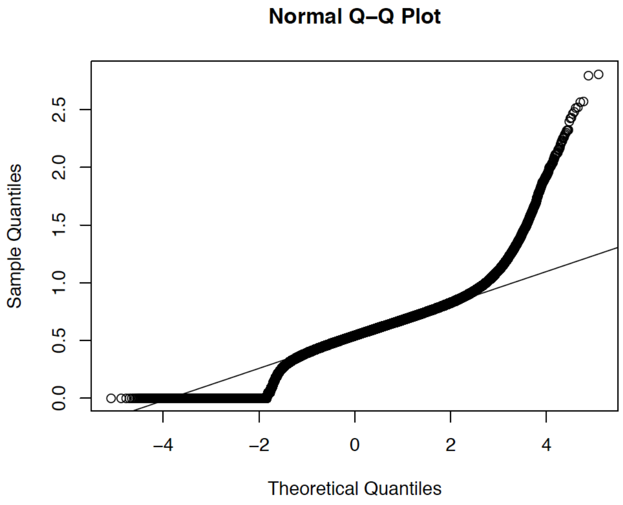

*Figure S 3. Q-Q plot of the log transformed signals for ChIP-seq data*

#### S3. BRCA1 survival analysis

Since BRCA1 demonstrates the highest number of rewired edges, as well as other significant regulatory changes between cancer and normal, we used the regulatory activity of BRCA1 as a marker for survival in acute myeloid leukemia (AML). We collected the cancer FPKM (Fragments Per Kilobase of transcript per Million mapped reads) expression data from 151 patients with AML from TCGA. In addition, we also collected the clinical information of each of these patients. Specifically, for deceased patients we used the number of days to death after diagnosis, while for alive patients, we used the days until the last follow up as a censored set of data for our survival analysis. We further compared the expression data for these cancer patients to a normal set of expression data from the GTex normal, whole blood cells. We scaled and took the difference between the cancer expression of each patient and the normal expression data in order to calculate the differential expression of each protein coding gene for each of the 151 patients. We extracted the network linking BRCA1 to over 2600 of the 20,000 protein coding genes, creating a binary vector of genes inside and outside of the BRCA1 network (represented by 1 and 0 respectively). A logistic regression was computed for each patient between this network vector and the set of differential expressions of each gene for each patient. The resulting coefficient for the regression was used as the regulatory activity for BRCA1 for a given patient. We then stratified the group of 151 patients by their high and low status of the BRCA1 regulatory activity. By stratifying the group in this way, we determine a statistically meaningful way of separating patients by quality of their prognosis.

#### S4. network centrality analysis

After selecting the optimal Lasso parameter $\lambda_{opt}$, we obtain the differential graphical model. In our case, we used $\lambda_{opt} = 0.2$. We then extracted all the connected edges in the graph $G$ and calculated the number of edges per node in $G$. We selected the top 10 nodes with the most edges as the network hubs and calculated the percentage of the network that includes each node in the 100 sub-sampled networks under $\lambda_{opt}$. In the end, we selected 8 out of the 10 nodes as consistent network hubs with inclusion rate of over 90 percent.

Table S1. List of Factors shared by GM12878 and K562 used in analysis

| List of Factors shared by GM12878 and K562 used in analysis | | | | | | | |
|---|---|---|---|---|---|---|---|
| ARID3A | CTCF | GABPA | MTA2 | NR2F1 | SIX5 | UBTF | ZSCAN29 |
| ARNT | CUX1 | GATAD2B | MTA3 | NRF1 | SKIL | USF1 | ZZZ3 |
| ATF2 | DPF2 | HCFC1 | MXI1 | PKNOX1 | SMAD1 | USF2 | H2AFZ |
| ATF7 | E2F4 | HDAC2 | NBN | POLR2A | SMAD5 | YBX1 | H3K27ac |
| BACH1 | E2F8 | HDGF | NFATC3 | POLR2AphosphoS2 | SMARCA5 | YY1 | H3K27me3 |
| BCLAF1 | E4F1 | IKZF1 | NFE2 | POLR2AphosphoS5 | SMC3 | ZBED1 | H3K36me3 |
| BHLHE40 | ELF1 | JUNB | NFIC | RAD51 | STAT5A | ZBTB33 | H3K4me1 |
| BMI1 | ELK1 | JUND | NFXL1 | RB1 | TAF1 | ZBTB40 | H3K4me2 |
| BRCA1 | EP300 | KDM1A | NFYA | RBBP5 | TARDBP | ZFP36 | H3K4me3 |
| CBX3 | ESRRA | LARP7 | NFYB | RCOR1 | TBL1XR1 | ZNF143 | H3K79me2 |
| CBX5 | ETS1 | MAX | NKRF | REST | TBP | ZNF24 | H3K9ac |
| CEBPB | ETV6 | MEF2A | NR2C1 | RFX5 | TCF12 | ZNF384 | DHS |
| CEBPZ | FOXK2 | MLLT1 | NR2C2 | SIN3A | TCF7 | ZNF592 | |
| CREM | | | | | | | |

#### S5. Pseudocode for model selection

### genome sampling and differential network estimation

for $s$ in 1:$S$:

Sample half of the columns from $\boldsymbol{X}^{(0)}$ as $\boldsymbol{X}^{(0),s}$

Sample half of the columns from $\boldsymbol{X}^{(1)}$ as $\boldsymbol{X}^{(1),s}$

for $m$ in $1{:}\,M$:

given $\lambda_m$, estimate differential network $\boldsymbol{N}_m^s$ with edge between TFs $i$ and $j$ if $(\psi_m^s)_{ij} \neq 0$

### genome sampling and differential network stability estimation

for $m$ in $1{:}\,M$:

$\widehat{D}(\Lambda_m) = \widehat{D}(1/\lambda_m) = 0$

for each possible edge $i, j$ $(i < j)$ in $(N_m^1, \cdots, N_m^s, \cdots, N_m^S)$ :

$\theta_{i,j}(\Lambda_m) = \frac{1}{S}\sum_{s=1}^{S} \psi_{i,j}^s(\Lambda_m)$

$\xi_{i,j}(\Lambda_m) = 2\theta_{i,j}(\Lambda_m)\{1 - \theta_{i,j}(\Lambda_m)\}$

$\widehat{D}(\Lambda_m) = \widehat{D}(\Lambda_m) + \xi_{i,j}(\Lambda_m)$

$\widehat{D}(\Lambda_m) = \frac{D(\Lambda_m)}{\binom{J}{2}}$

### monotone network stability

for $m$ in $1{:}\,M$:

$\overline{D}(\Lambda_m) = sup_{0 \ll t \ll \lambda_m} \widehat{D}(\Lambda_m)$

### optimal $\lambda$ selection

$\widehat{\Lambda}_{opt} = sup\{\Lambda{:}\,\overline{D}(\Lambda) \leq \beta\}$
